# An introduced parasitoid facilitates host range expansion of a resident parasitoid

**DOI:** 10.64898/2025.12.22.696021

**Authors:** Jessie Moon, Jessica L. Fraser, Paul K. Abram

## Abstract

Invasive species can cause maladaptive behaviours in native consumers, with evolution of behavioural or physiological traits thought to be the main way out of these so-called ‘evolutionary traps’. However, shorter-term ecological processes resulting from biological invasions could provide other ways to escape evolutionary traps. *Asobara* cf. *rufescens*, a resident (i.e., possibly native or previously introduced) parasitoid of drosophilid fly larvae in North America, can rarely produce offspring on the invasive fruit pest *Drosophila suzukii* in the laboratory, but in often emerges from field-collected *D. suzukii*. We hypothesized that the successful development of *A.* cf. *rufescens* in *D. suzukii* in the field is facilitated by *Leptopilina japonica*, a recently introduced parasitoid of *D. suzukii*. In laboratory experiments, we found that *A.* cf. *rufescens* had more than 30-fold higher offspring emergence from *D. suzukii* when *L. japonica* was also present. This facilitation occurred most frequently when *A.* cf. *rufescens* parasitized *D. suzukii* after *L. japonica*, possibly because parasitism by *L. japonica* destroyed the hosts’ cellular immunity that would otherwise prevent *A.* cf. *rufescens* development. The recent arrival of *L. japonica* may have partially rescued *A.* cf. *rufescens* from the evolutionary trap set by *D. suzukii* and, consequently, resulted in an expansion of its host range.

## Introduction

Native herbivores, predators and parasites may sometimes suffer negative fitness consequences when they attempt to use invasive species as prey or hosts. These ‘evolutionary traps’ result from a disconnect between the cues used by consumers to identify resources and the resulting suitability of the invasive species as a resource [1]. Native species can escape evolutionary traps by evolving morphological, behavioural or physiological traits that prevent them from consuming the unsuitable invasive species or allow them to more successfully exploit the invasive species as a host [2–4]. These evolutionary processes are expected to take decades, at least, to play out. There may also, however, be shorter-term ecological processes that could allow native species to escape evolutionary traps.

Parasitoid wasps are a hyper-diverse group of insects whose offspring develop on or inside the bodies of other insects and kill them as a result. Some parasitoids make ‘paradoxical’ egg-laying decisions: they deposit eggs on or into hosts that are unsuitable for their offspring’s development [5–10]. Interestingly, facilitation by one parasitoid species can allow parasitoid offspring of another species to complete development in an otherwise unsuitable host, ‘rescuing’ them from almost certain death. Such facilitation can result from kleptoparasitism (resource-stealing by the facilitated parasitoid) [5,11,12,13] or hyperparasitism (parasitism of the facilitating parasitoid by the facilitated parasitoid) [14]. Some kleptoparasitoids, whose offspring’s development would otherwise be thwarted by host immune responses, can take advantage of the facilitating parasitoid’s destruction of the hosts’ cellular immunity, kill the facilitating parasitoid’s offspring through larval competition, and steal the host resource [5,11,12,13]. If parasitism by the facilitating species is common enough, parasitism of otherwise unsuitable hosts by the facilitated species could be favoured by natural selection [5]. This would be expected to evolve in ecological systems where the facilitating and facilitated parasitoid species have been in sympatry for relatively long time periods [12].

However, in some cases parasitism of unsuitable invasive hosts could simply be a maladaptive behaviour from which parasitoids can be incidentally ‘rescued’ by other, facilitating parasitoids. One laboratory study showed that a native parasitoid could kleptoparasitize an invasive host whose immune systems had been subdued by a native generalist parasitoid [13]. In another laboratory study, a native parasitoid species could not successfully parasitize an invasive pest’s eggs, but could hyperparasitize eggs parasitized by a more recently introduced, specialized parasitoid species during a short (∼24h) window of the introduced parasitoid’s development [14]. In the latter case, the arrival of the introduced parasitoid was hypothesized to be an ‘invasional lifeline’ for the native parasitoid, allowing the native parasitoid to partially escape the evolutionary trap posed by the invasive host insect [14]. However, in neither case are there reports of how often these native parasitoids parasitize the invasive pests in the field, so it is unclear whether these types of facilitative interactions are lab artefacts or whether they may actually allow host range expansions to occur under natural conditions.

Here we report a likely example of an invasional lifeline whose discovery was built on field-based natural history observations made over a period of ∼15 years in British Columbia, Canada [15–21]. The vinegar fly *Drosophila suzukii*, native to Asia, first invaded North America and Europe in 2008–2009 and became one of the most serious agricultural pests of soft fruit crops [22]. Laboratory studies found that some native and long-present cosmopolitan (hereafter, ‘resident’) parasitoids in invaded areas would readily lay their eggs in *D. suzukii* larvae, but that the immune system of the host would prevent parasitoid development, suggesting that *D. suzukii* represents an evolutionary trap for these species [4,9,23].

Beginning in 2021, the resident parasitoid *Asobara* cf. *rufescens* was found to commonly emerge from field-collected *D. suzukii* puparia extracted from soil and rotting fruit in British Columbia [17,19,21] (Figure 1). In these studies, *A.* cf. *rufescens* emerged from ground-collected *D. suzukii* at all sites in every year, and represented up to 68% of all emerging parasitoids. This was surprising, given that field studies using several collecting techniques in the same region from 2009–2011 never reared *A.* cf. *rufescens* from *D. suzukii*, but did rear it from other drosophilid species [15,20]. Likewise, previous laboratory observations showed that *A.* cf. *rufescens* and several other closely related *Asobara* species readily attack *D. suzukii* but rarely or never produce offspring [16,23,24]. This led us to ask what may have changed to allow *A.* cf. *rufescens* to apparently expand its realized host range to include *D. suzukii* in western North America (Figure 1).

**Figure 1.**
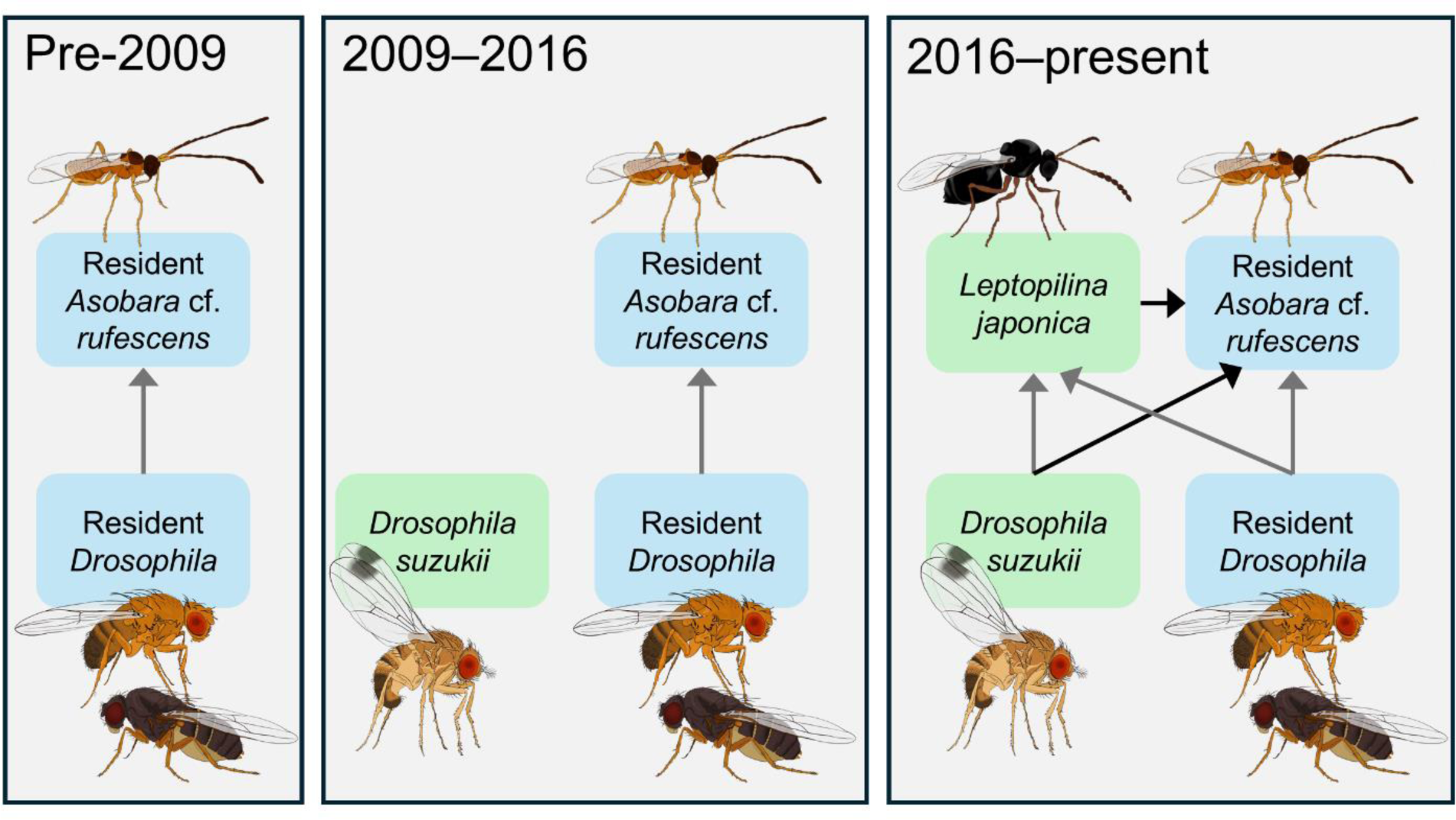
Simplified *Drosophila*-larval parasitoid food webs in the study region over time. Arrows show food web links; arrowheads point in the direction of energy flow. Black arrows in the right-most panel show the ecological interaction that this study addresses (*A.* cf. *rufescens* developing in *D. suzukii*) and the hypothesized facilitation of *A.* cf. *rufescens* by *L. japonica*. For simplicity, other resident and introduced parasitoid species are not shown (see [21] for details).

Other *Asobara* species are known or suspected to be kleptoparasitic [5,24]. We therefore reasoned that the change leading to the host range expansion of *A.* cf. *rufescens* could be the recent establishment of parasitoid wasps from Asia that are well adapted to *D. suzukii*, *Leptopilina japonica* and *Ganaspis kimorum* [16,18,20] (Figure 1). Specifically, we hypothesized that parasitism by one or both of the recently introduced parasitoids facilitates the development of *A.* cf. *rufescens* inside *D. suzukii* larvae. Here we report on laboratory experiments with *A.* cf. *rufescens* and the most common of the two introduced parasitoids, *L. japonica*, that support this hypothesis.

## Materials and Methods

### Insect colonies

*Drosophila suzukii* colonies, established from populations collected in Agassiz and Vancouver (British Columbia, Canada), were maintained on artificial diet (Carolina Drosophila 4-24 medium, Ayva Educational Solutions Ltd., Canada) in plastic vials (diameter: 2 cm; height: 9 cm, Diamed, Canada) under laboratory conditions (22–24°C, 16:8 light:dark cycle; 55–65% relative humidity) for less than a year before each experiment was conducted. Flies used in experiments were first transferred to vials containing lower densities of mixed-sex flies (<20 individuals) within 72 h of emergence. These groups of flies were transferred into vials with fresh medium twice weekly until their use in experiments.

Colonies of *L. japonica* and *A.* cf. *rufescens* used in Experiment 1 were collected in Agassiz, British Columbia, Canada in 2022 and 2023. Both wasps were reared on *D. suzukii* and were provided with liquid honey for nutrition *ad libitum*. *Asobara* cf. *rufescens* could not be reared on *D. suzukii* alone, and so was reared on *D. suzukii* in the presence of *L. japonica*. Parasitoid adults used in experiments were first transferred to vials in smaller groups (<20 individuals) within 72h of emergence. These vials contained cotton moistened with water, and liquid honey smeared on the underside of the vial plug.

All experiments were conducted at 22–24°C, a 16:8 light: dark cycle, and 55–65% relative humidity. The taxonomic identity of voucher specimens were confirmed by experts and additional vouchers are stored at the Agassiz Research and Development Centre, Agassiz, BC, Canada.

### Experiment 1: Testing for facilitation

To test whether the parasitism of *D. suzukii* by *L. japonica* facilitates *A.* cf. *rufescens*, we exposed *D. suzukii* to each of the two parasitoid species alone or in combination. Test arenas were large Drosophila vials (diameter: 2.8 cm; height: 9.5 cm) (Diamed, Canada) containing one grocery store-bought raspberry placed on top of crumpled paper towel (to absorb liquid). A pinch of baker’s yeast was added and a small quantity (∼4 mL) of Drosophila medium was smeared on the side of the vial to supply nutrition to adult flies. Depending on the availability of adequately aged flies, between 5 and 10 female, mated *D. suzukii* (7–21 days old) were added to each vial for 3–4 days to lay eggs that would develop into larvae. Female parasitoids (1–7 days old, mated) were then added to vials for a period of 7 days in one of the following three combinations: (1) Two female *A. cf. rufescens*; (2) Two female *L. japonica*; (3) Two females of each species. Liquid honey was smeared *ad libitum* on the underside of the vial plugs to provide nutrition to wasps. After parasitoids were removed, vials were incubated under standard conditions (see above) for at least 30 days, and adult *D. suzukii* were removed as they emerged. The number of emerging parasitoids of each species from each vial was then counted. Nine replicates of each of the three treatments were conducted (=27 vials total) over 7 temporal blocks of 1-2 full replications each.

### Experiment 2: The effect of the order of parasitism on facilitation

To test whether the facilitation of *A.* cf. *rufescens* by *L. japonica* depended on the order in which each species parasitized *D. suzukii*, we conducted an experiment varying the order of introduction of each parasitoid species to vials containing *D. suzukii* larvae. Conditions were more tightly controlled than in Experiment 1 (single parasitoid individuals, much shorter time intervals of 24h) to test whether *A.* cf. *rufescens* could develop in larvae parasitized by *D. suzukii* when *L. japonica* were still in the egg or early-instar larval stages. Each test arena was made by placing 8 female *D. suzukii* (aged 7–14 days) in a plastic vial (diameter: 2 cm; height: 9 cm) capped with a cellulose plug containing ∼4 mL of diet to allow the flies to lay eggs. After 24 hours, flies were removed, and a 3.0 cm-long cotton dental wick was placed into the media to absorb excess liquid. Liquid honey was smeared on the underside of the vial plug. The vials were then divided into six treatments determining the timing and order of female parasitoid introductions: (1) One *A. rufescens* for 24h; (2) One *L. japonica* for 24h; (3) One *L. japonica* and one *A.* cf. *rufescens* added together for 24h; (4) *L. japonica* for 24h, then no parasitoids for 24h, then *A.* cf. *rufescens* for 24h; (5) *A.* cf. *rufescens* for 24h, then no parasitoids for 24h, then *L. japonica* for 24h; (6) no parasitoids (included as a control to compare the rate of encapsulation responses; see below). Parasitoids were 2–7 days old when introduced to vials. Five temporal blocks of 3-4 full replications each were conducted, for a total of 17 replicates per treatment. Vials were incubated for 30 days under standard rearing conditions and emerging *D. suzukii* and parasitoids were preserved in 95% EtOH until they were counted. In addition, to confirm that *A.* cf. *rufescens* was unsuccessfully parasitizing *D. suzukii* in the absence of *L. japonica* (with the no-parasitoid treatment as a reference), emerging adult *D. suzukii* were also compressed between two glass slides under 40× magnification to count the number of black melanic capsules within their bodies, which serve as evidence of failed parasitism due to the host larva’s encapsulation response [25]. Encapsulations were identified according to consistent criteria (oblong shape, opaque black coloration, relatively large size). Black capsules were rarely found in *D. suzukii* larvae not exposed to parasitoids, indicating that other processes (e.g., injury) could generate similar structures. However, comparison of the presence of black capsules between the control (no parasitoids) and the other (parasitoid-exposed) treatments provided an approximation of the relative incidence of failed parasitism among treatments.

### Data analysis

All statistical analyses were conducted in R version 4.4.1 [26]. For Experiment 1, we ran two negative binomial generalized linear models (GLMs) with the number of parasitoids of each species emerging as response variable and treatment as the predictor variable. For Experiment 2, where the shorter duration of host exposure to parasitoids likely led to a greater number of replicates in which no parasitoids emerged, we used zero-inflated negative binomial GLMs implemented with glmmTMB [27] to test how the number of emerging parasitoids of each species and the number of black capsules in emerging adult flies responded to different treatments. For all response variables, multiple comparisons among treatments for Experiment 2 were done with the Tukey method, implemented with emmeans [28]. Type II likelihood ratio tests were used to test the statistical significance of the treatment factor for both experiments. Comparisons of the number of offspring emerging for each of the two parasitoid species were only done among treatments which contained that species.

## Results

### Experiment 1: Testing for facilitation

In the absence of parasitism by *L. japonica*, *A.* cf. *rufescens* offered *D. suzukii* larvae over a period of 7 days rarely produced offspring (Figure 2A) (∼0.44 offspring per replicate). In contrast, ∼34 times more *A.* cf. *rufescens* offspring emerged (∼15.11 offspring per replicate) when *L. japonica* was also present (χ^2^1,16 = 41.82, p < 0.0001) (Figure 2A).

**Figure 2.**
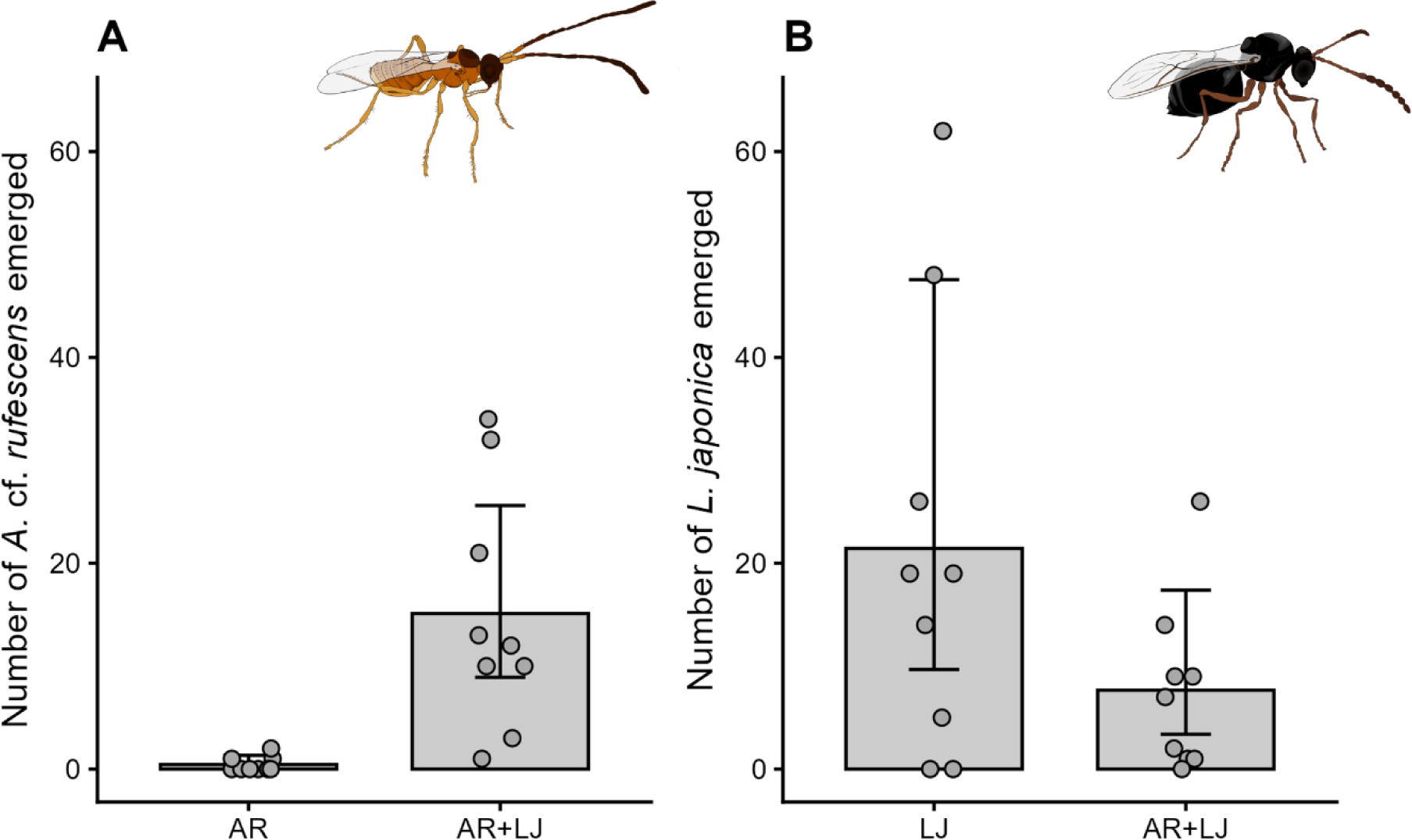
The mean (model estimate ± 95% CI) number of (A) *Asobara* cf. *rufescens* (AR) and (B) *Leptopilina japonica* (LJ) emerging when alone (AR, LJ) or both species together (AR + LJ) were offered *Drosophila suzukii* larvae.

Approximately 60% fewer *L. japonica* emerged when *L. japonica* was offered *D. suzukii* larvae in the presence of *A.* cf. *rufescens*, compared to when *A.* cf. *rufescens* was not present, although this difference was only marginally statistically significant (χ^2^1,16 = 3.01, p = 0.083) (Figure 2B).

### Experiment 2: The effect of the order of parasitism on facilitation

Different combinations and ordering of shorter-term (24 h) parasitoid exposures to *D. suzukii* resulted in differences in the number of emerging *A.* cf. *rufescens* (χ^2^3,62 = 7.76, p = 0.051) and *L. japonica* (χ^2^3,62 = 21.22, p < 0.0001) (Figure 3). Zero *A.* cf. *rufescens* emerged when offered *D*. *suzukii* larvae in the absence of *L. japonica*, and only a single individual emerged when *A.* cf. *rufescens* was offered host larvae before *L. japonica*. The most *A.* cf. *rufescens* offspring emerged when *L. japonica* was offered host larvae before *A.* cf. *rufescens*, and an intermediate number emerged when *D. suzukii* was exposed to both parasitoids during the same 24 h period (Figure 3). The number of emerging *L. japonica* was lowest when *L. japonica* was added after *A.* cf. *rufescens* (Figure 3).

**Figure 3.**
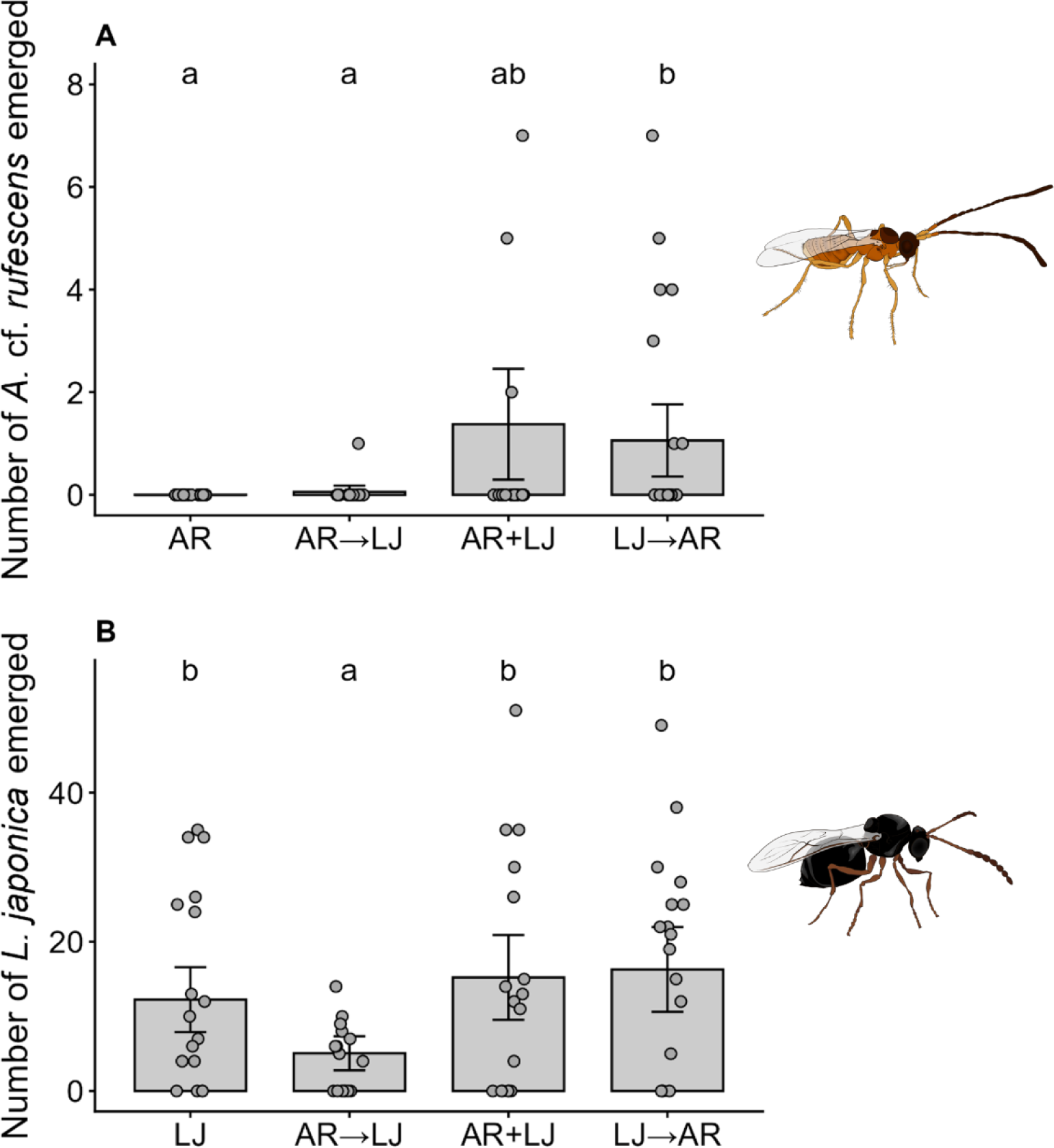
The mean (model estimate ± 95% CI) number of (A) *Asobara* cf. *rufescens* (AR) and (B) *Leptopilina japonica* (LJ) emerging when offered *Drosophila suzukii* larvae alone (AR – *A*. cf. *rufescens* alone; LJ – *L*. *japonica* alone) or in combination (+ indicates both species were added at the same time; → indicates that one species was removed, then the other species was added 24h later). Means not labeled with the same letter are statistically different (p < 0.05; Tukey-adjusted multiple comparisons).

The number of emerging flies bearing black capsules varied among treatments (χ^2^5,94 = 19.61, p < 0.01). The only treatment where more *D. suzukii* containing black capsules were observed compared to the control without parasitoids was when *A.* cf. *rufescens* was exposed to *D. suzukii* without *L. japonica* (Online Supplementary Material; Figure S1).

## Discussion

Parasitism of *D. suzukii* by the recently introduced parasitoid *L. japonica* clearly facilitated the development of the resident parasitoid *A.* cf. *rufescens* in the laboratory, especially when parasitism by *A.* cf. *rufescens* took place after *L. japonica*. Given that *L. japonica* is now by far the most common larval parasitoid of *D. suzukii* in the field [18–21,29], these results suggest that the unexpected emergence of *A.* cf. *rufescens* from *D. suzukii* is largely a result of facilitation by *L. japonica*. Further studies would be needed to determine whether the other much less common introduced larval parasitoid, *G. kimorum*, may also be able to facilitate *A.* cf. *rufescens* development.

The expansion of the host range of *A.* cf. *rufescens* to include *L. japonica-*parasitized *D. suzukii* has the potential to considerably increase overall host availability for *A.* cf. *rufescens* in nature. Indeed, *D. suzukii* can be highly abundant, and an average of ∼25% (up to 40%) of *D. suzukii* larvae in fruit that have dropped from plants on the ground may be parasitized by *L. japonica* [21]. Strikingly, *A.* cf. *rufescens* has recently become one of the main parasitoids emerging from ground-collected *D. suzukii* [17,19,21]. Thus, invasive species that are highly abundant may at first be potent evolutionary traps for resident parasitoids, but can also create opportunities in terms of resource availability for consumer species that are able to escape the trap. Researchers in other areas of the world where *L. japonica* has recently been introduced [20,29] could now investigate whether other resident parasitoid species have been rescued from the evolutionary trap set by *D. suzukii*’s invasion.

The mechanism by which *L. japonica* facilitates *A.* cf. *rufescens* has yet to be determined and is outside the scope of the current study. The most likely candidate explanation, however, is that *A.* cf. *rufescens* acts as a kleptoparasitoid in *D. suzukii* parasitized by *L. japonica*. Other *Leptopilina* species inject virus-like particles into their hosts that destroy their cellular immunity and prevent them from encapsulating the parasitoid’s eggs [30,31]. *Asobara* species are known to lack these particles [5,32], and in our experiments the highest rates of encapsulation were observed when *A.* cf. *rufescens* was offered *D. suzukii* in the absence of *L. japonica*, confirming that it did unsuccessfully attack *D. suzukii* in the absence of *L. japonica*. Thus, the destruction of host immunity by *L. japonica* may ‘open the door’ to *A.* cf. *rufescens* – which develops more rapidly [19] and thus may have a competitive advantage – stealing the host resource from *L. japonica*. Hyperparasitism is an unlikely mechanism to explain the observed facilitation given that *L. japonica* would still be in the egg and early larval stage (and thus too small to hyperparasitize) during the time window where *A. rufescens* was offered hosts. Further research is needed to uncover the details of how this facilitation occurs. In addition, applying molecular diagnostic tools [20] to field-collected *D. suzukii* larvae will help to define patterns of unsuccessful parasitism and multiparasitism among parasitoids of *D. suzukii*.

There has been considerable international interest in *L. japonica* as a biological pest control agent for *D. suzukii* [29]. Our study raises the question of what consequences its facilitation of a resident parasitoid species may have for biological control of *D. suzukii*. *Asobara* cf. *rufescens* appears to be parasitizing *D. suzukii* that would have already died from parasitism by *L. japonica*, and the number of hosts emerging from different treatments in Experiment 2 do not suggest that *Asobara* kill significant numbers of *D. suzukii* with unsuccessful parasitism attempts (Online Supplementary Material; Figure S2). Because this facilitative interaction results in the death of *L. japonica*, it is likely to be counterproductive for biological control of *D. suzukii*. From a population dynamics perspective the outcomes may be similar to hyperparasitism [33,34].

Several authors have hypothesized that parasitoids could escape an evolutionary trap by evolving the ability to develop in unsuitable host species, resulting in an expansion of their host range [4,8,9,35,36]. This represents a rather counterintuitive process of a behaviour that is maladaptive at the individual level eventually leading to broadening of host range at the species level. Our study – which was inspired by field-based observations of a recent host range expansion – adds to the evidence that the evolution by natural selection is, in fact, not the only way for a parasitoid wasp species to expand its host range to include previously unsuitable host species. Shorter-term ecological processes, like biological invasions by other parasitoids, may occasionally be able to provide a faster way out of evolutionary traps *en route* to host range expansions.

## Acknowledgements

We thank Warren Wong, Jiamin Bai, Fina VanderPloeg, Jason Thiessen, Emily Grove, Madison Khashan, and Ethan Buchanan for help in the field and laboratory. We thank Matt Buffington and Robert Kula (USDA-ARS/Smithsonian Museum of Natural History, Washington, DC, USA) for taxonomic support and collaboration, and Xingeng Wang (USDA-ARS) for helpful discussions and methodological advice. Thanks to Emily Grove for creating the insect artwork. Thanks to Jacques Brodeur for helpful comments on an earlier version of the manuscript.

