## Supplementary material for "An introduced parasitoid facilitates host range expansion of a resident parasitoid": Online Supplementary Material

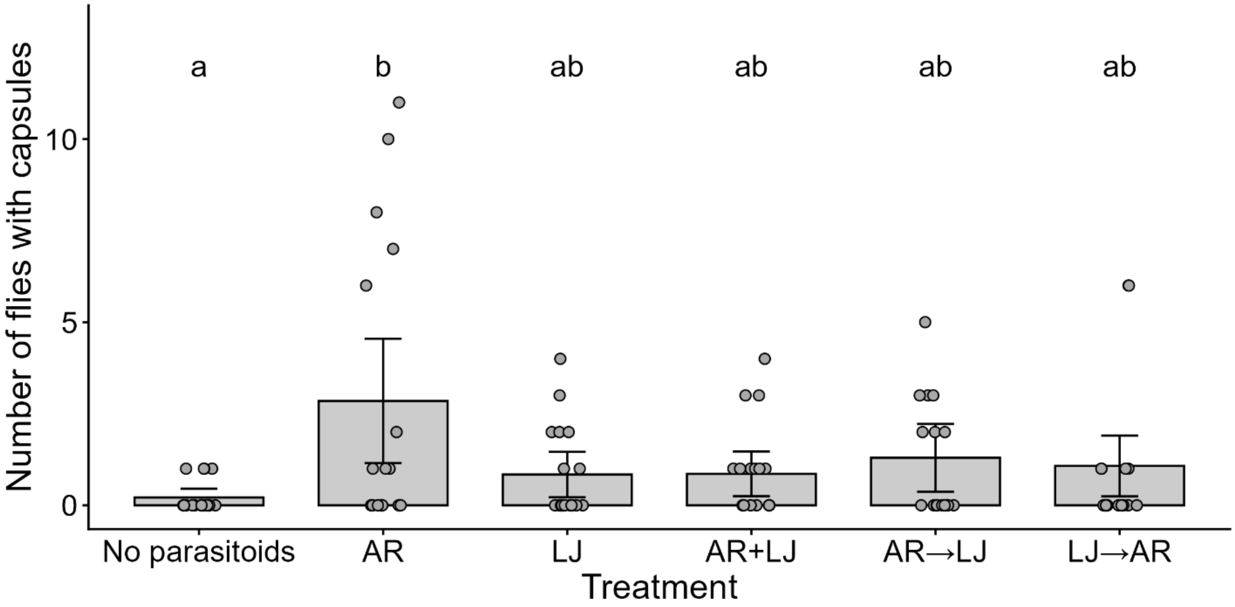


**Figure S1**. The mean (model estimate ± 95% CI) number of black capsules observed in emerging *D. suzukii* adults when they were offered as larvae to no parasitoids, individual parasitoid species (AR – *A*. cf. *rufescens* alone; LJ – *L. japonica* alone), or the two parasitoid species in combination (+ indicates both species were added at the same time; → indicates that one species was removed, then the other species was added 24h later). Means not labeled with the same letter are statistically different ($p < 0.05$; Tukey-adjusted multiple comparisons).

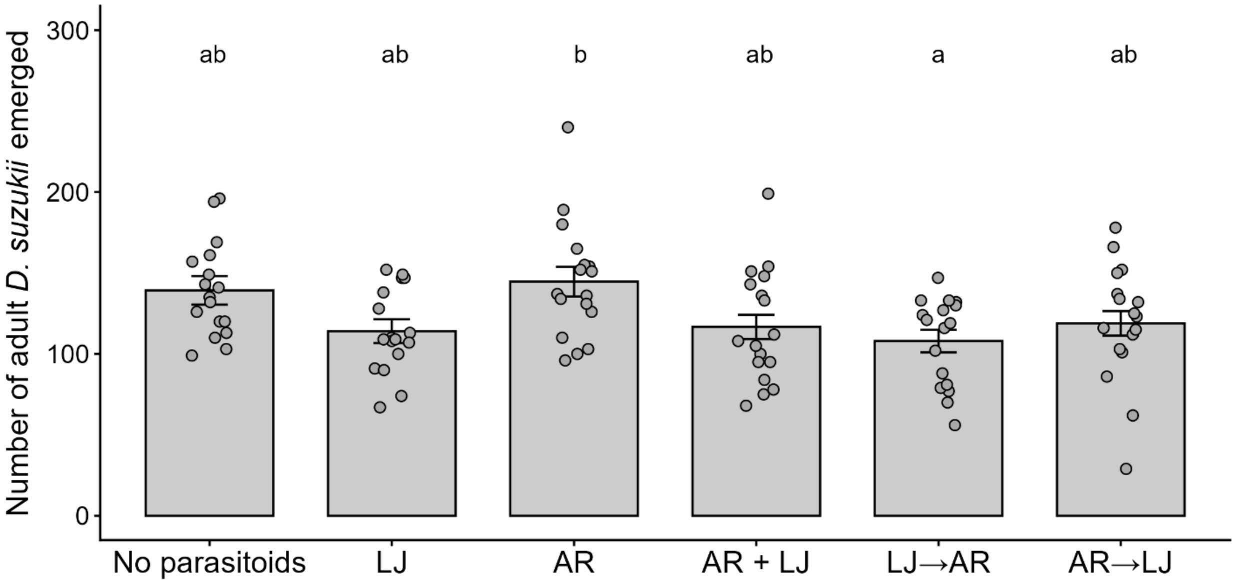


**Figure S2.** The mean (model estimate ± 95% CI) number emerging *D. suzukii* adults when larvae were offered to no parasitoids, individual parasitoids (AR – *A*. cf. *rufescens* alone; LJ – *L. japonica* alone), or the two parasitoid species in combination (+ indicates both species were added at the same time; → indicates that one species was removed, then the other species was added 24h later). Means not labeled with the same letter are statistically different ($p < 0.05$; Tukey-adjusted multiple comparisons). The number of emerging *D. suzukii* varied among treatments ($\chi^2_{5,96} = 17.07$, $p < 0.01$), but there were no treatments where the presence of parasitoids reduced the number of emerging *D. suzukii* below those observed in the absence of parasitoids (Figure S2). Fewer *D. suzukii* emerged when *L. japonica* was offered hosts before *A.* cf. *rufescens* than when *A.* cf. *rufescens* was offered hosts alone. There were no clear differences in *D. suzukii* emergence among any of the other treatments.
